## Supplementary figures and images for "Discarded sequencing reads uncover natural variation in pest resistance in *Thlaspi arvense*"

### S1 Fig

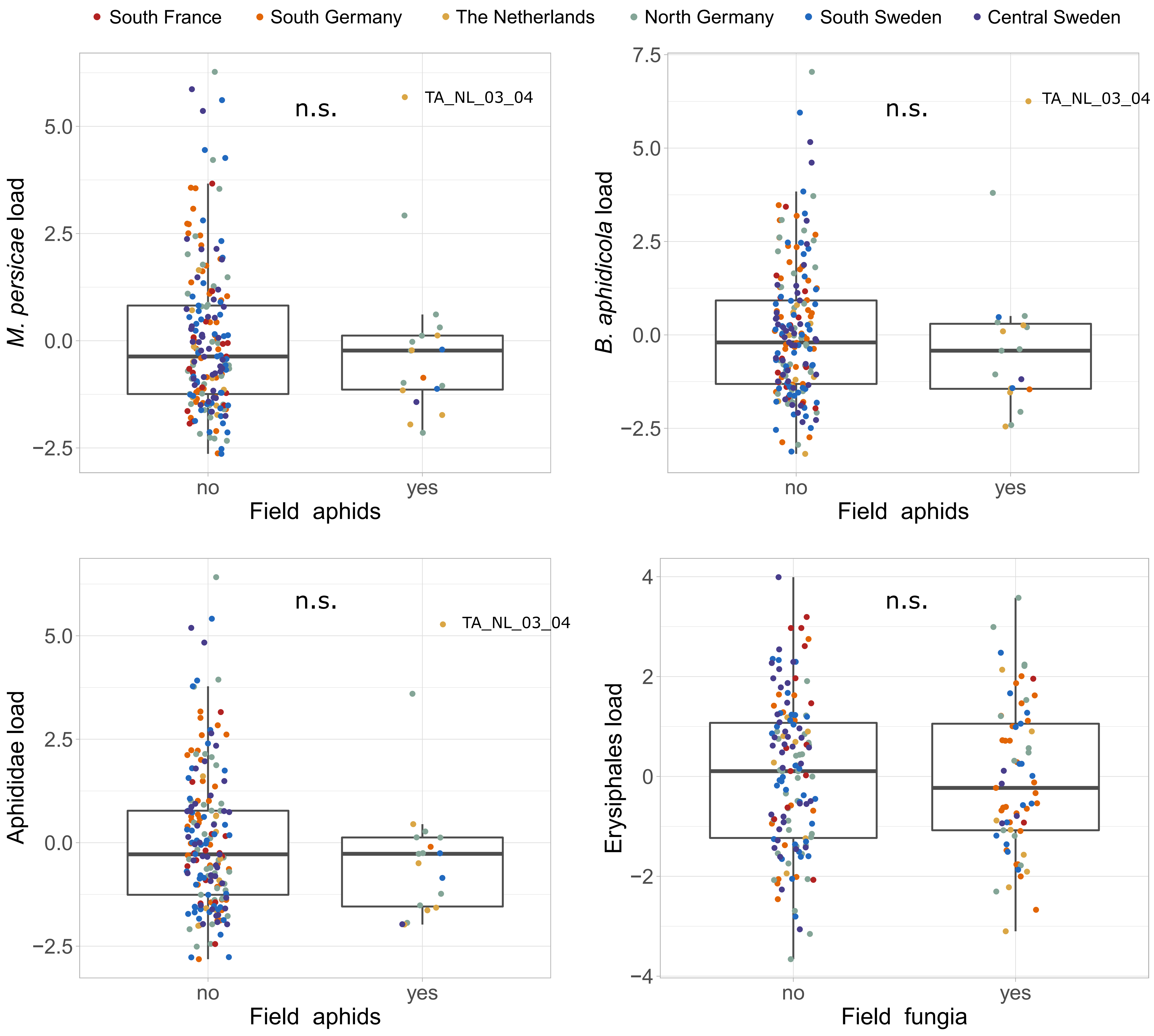

### S2 Fig

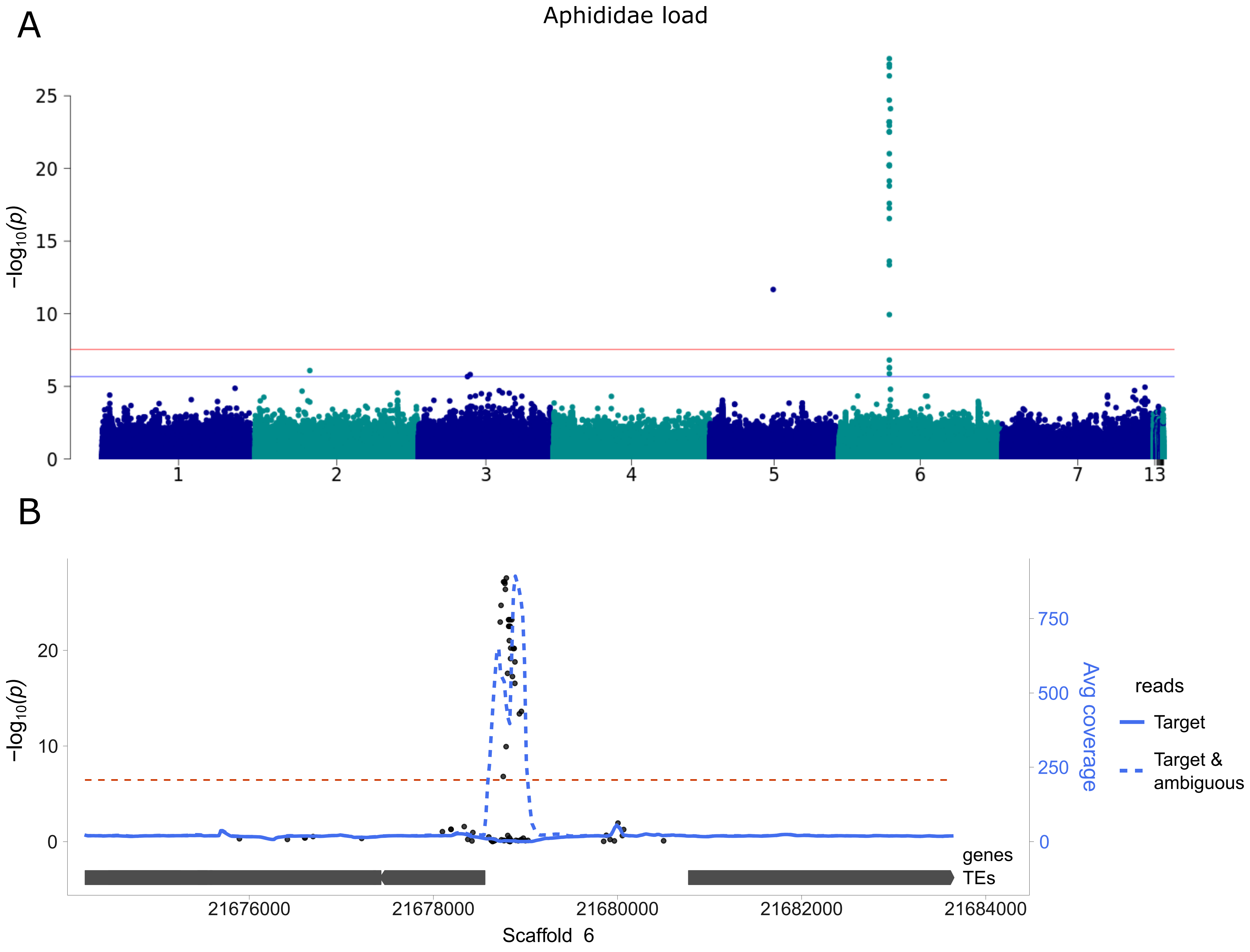

### S3 Fig

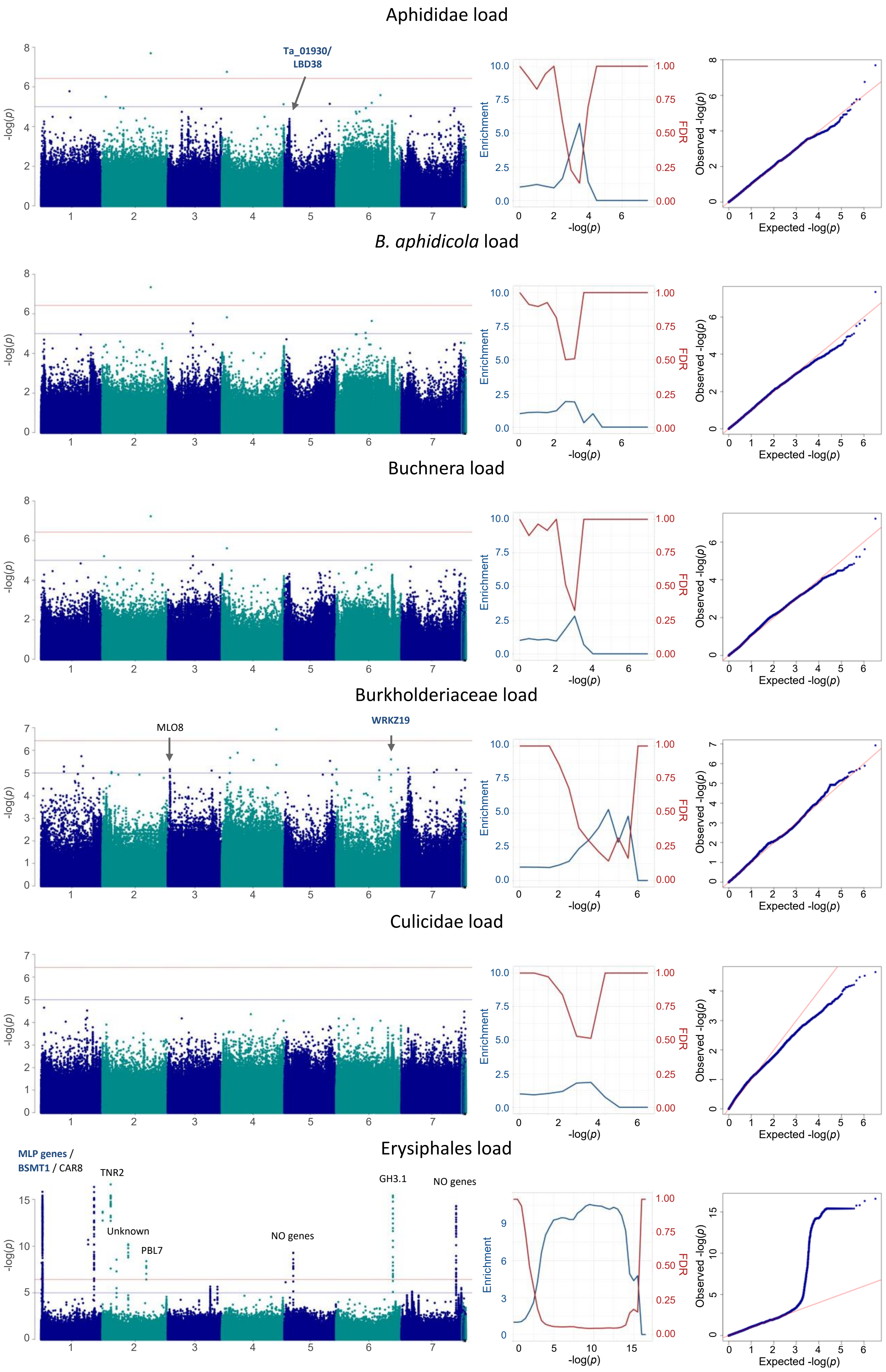

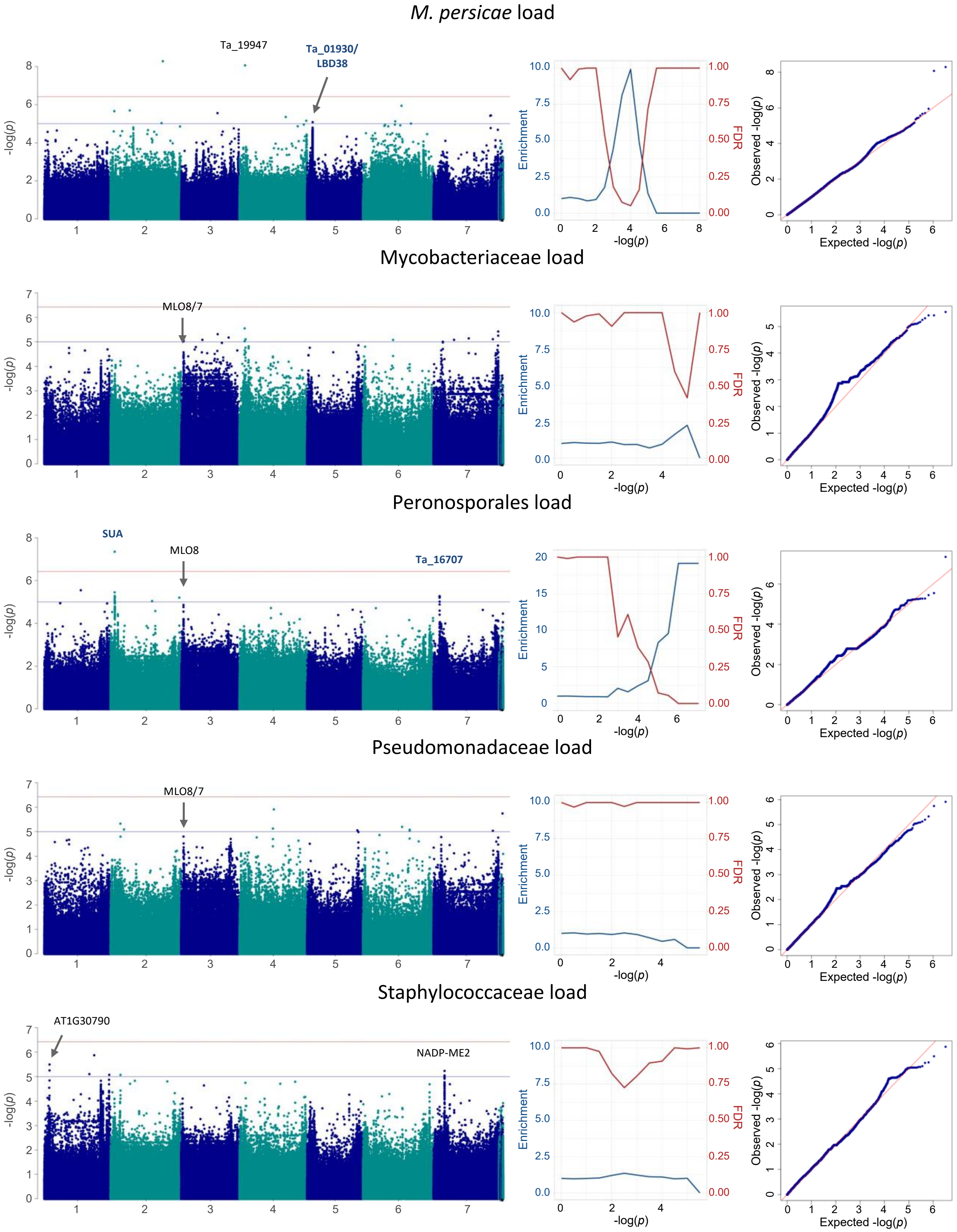

### S4 Fig

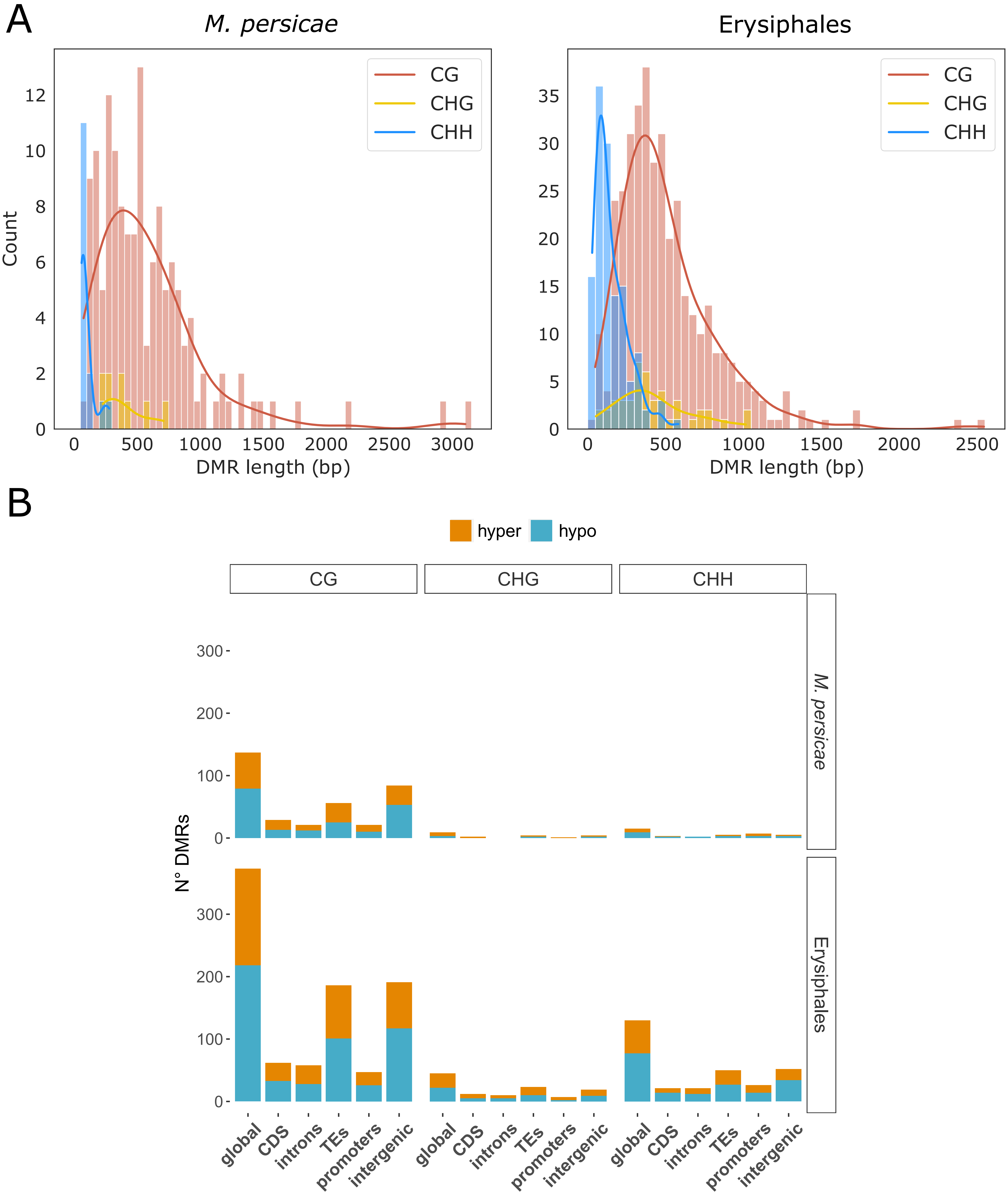

### S5 Fig

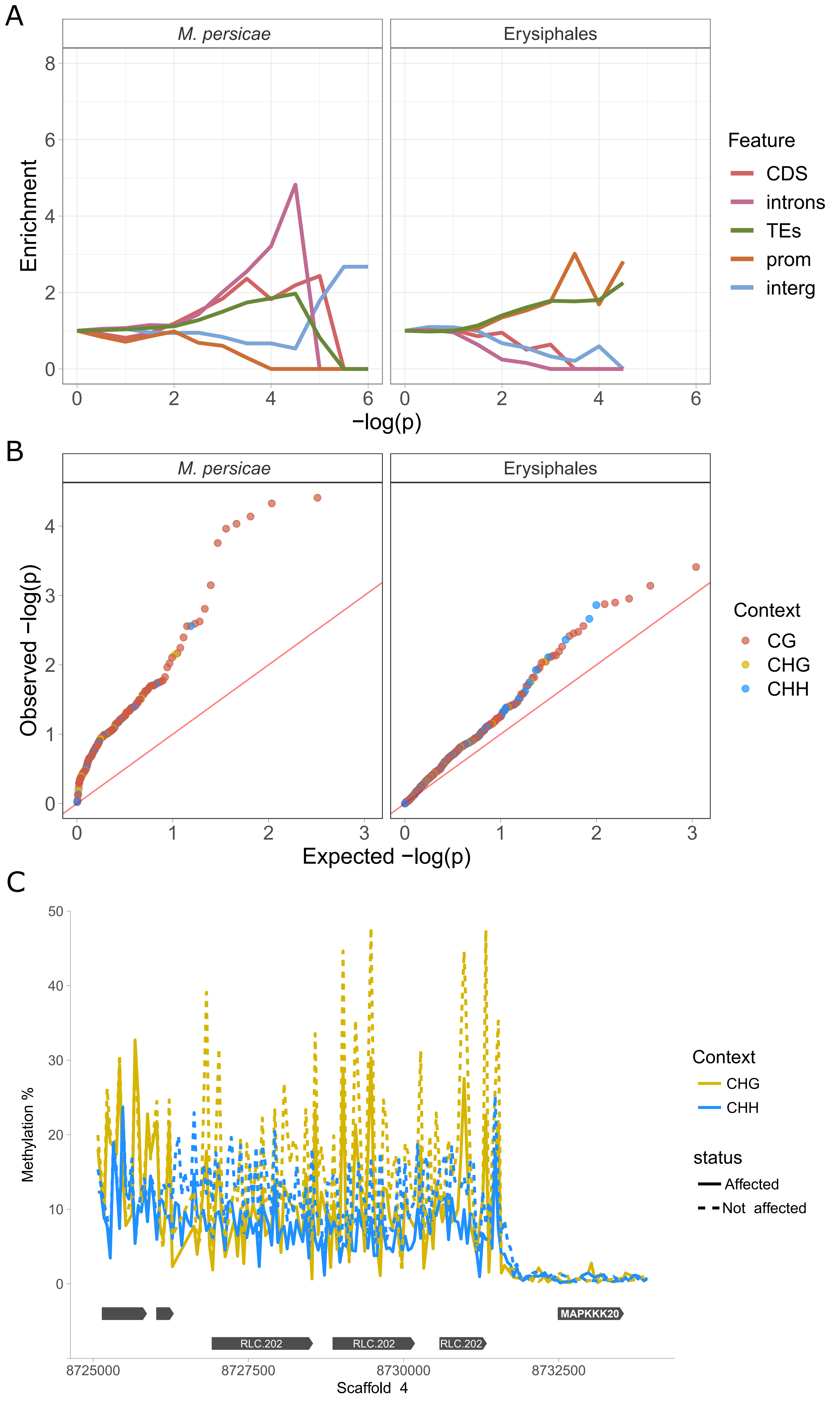
